# Using machine learning to predict antimicrobial minimum inhibitory concentrations and associated genomic features for nontyphoidal *Salmonella*

**DOI:** 10.1101/380782

**Authors:** Marcus Nguyen, S. Wesley Long, Patrick F. McDermott, Randall J. Olsen, Robert Olson, Rick L. Stevens, Gregory H. Tyson, Shaohua Zhao, James J. Davis

**Affiliations:** University of Chicago Consortium for Advanced Science and Engineering, University of Chicago, Chicago, Illinois, 60637, USA; Computing, Environment and Life Sciences, Argonne National Laboratory, Argonne IL, 60439, USA; Center for Molecular and Translational Human Infectious Diseases Research, Department of Pathology and Genomic Medicine, Houston Methodist Research Institute and Houston Methodist Hospital, Houston, Texas, 77030, USA; Department of Pathology and Laboratory Medicine, Weill Cornell Medical College, New York, New York, 10065, USA; Food and Drug Administration, Center for Veterinary Medicine, Office of Research, Laurel, MD 20708, USA; University of Chicago, Department of Computer Science, Chicago, IL, 60439, USA

**Keywords:** Machine learning, Deep learning, Antimicrobial susceptibility testing, Genome sequencing, Diagnostics

## Abstract

Nontyphoidal *Salmonella* species are the leading bacterial cause of food-borne disease in the United States. Whole genome sequences and paired antimicrobial susceptibility data are available for *Salmonella* strains because of surveillance efforts from public health agencies. In this study, a collection of 5,278 nontyphoidal *Salmonella* genomes, collected over 15 years in the United States, were used to generate XGBoost-based machine learning models for predicting minimum inhibitory concentrations (MICs) for 15 antibiotics. The MIC prediction models have average accuracies between 95-96% within ± 1 two-fold dilution factor and can predict MICs with no *a priori* information about the underlying gene content or resistance phenotypes of the strains. By selecting diverse genomes for training sets, we show that highly accurate MIC prediction models can be generated with fewer than 500 genomes. We also show that our approach for predicting MICs is stable over time despite annual fluctuations in antimicrobial resistance gene content in the sampled genomes. Finally, using feature selection, we explore the important genomic regions identified by the models for predicting MICs. To date, this is one of the largest MIC modeling studies to be published. Our strategy for developing whole genome sequence-based models for surveillance and clinical diagnostics can be readily applied to other important human pathogens.

## Introduction

Nontyphoidal *Salmonella* species are the leading bacterial cause of food-borne disease in the United States[1, 2], causing over one million illnesses per year[3] and an estimated 80 million illnesses annually world-wide[4]. Nontyphoidal *Salmonella* causes acute gastroenteritis and is usually contracted via fecal contamination of food sources[5]. Although these infections are usually self-limiting and typically do not require antibiotic treatment[6], severe infections can occur[7]. Antimicrobial resistance (AMR) is prevalent in *Salmonella* isolates and infections caused by highly antimicrobial resistant *Salmonella* strains result in worse outcomes than infections caused by susceptible strains[8–11].

In 1996, the National Antimicrobial Resistance Monitoring System (NARMS) was established as a collaboration between the United States Centers for Disease Control and Prevention (CDC), U.S. Food and Drug Administration (FDA), U.S. Department of Agriculture (USDA), and state and local health departments. A primary goal of NARMS is to monitor antimicrobial resistance in *Salmonella* and other food-borne bacteria, including *Campylobacter, Escherichia* and *Enterococcus*[12]. The data collected by NARMS is used to inform public health decisions aimed at identifying contaminated food sources and reducing the spread of AMR through enhanced stewardship. In recent years, NARMS has adopted whole genome sequencing (WGS) as a routine monitoring tool. The WGS data are used to determine the source of outbreak strains, the virulence factor and AMR genes carried by each strain. As a result, a large collection of bacterial whole genome sequences with extensive metadata is available for downstream research efforts[13].

Whole genome sequencing is now routinely used for public health surveillance and to guide diagnostic and patient care descisions[14–18]. For routine surveillance, WGS provides the highest possible resolution for individuating traits in bacteria, assessing phylogenetic relationships, conducting outbreak investigations, and predicting virulence and epidemicity. From the clinical perspective, rapid diagnostics are key to improving patient care. For a conventional microbiology laboratory diagnosis, the total time for organism growth, isolation, taxonomic identification, and antimicrobial minimum inhibitory concentration (MIC) determination may exceed 36 hours for relatively fast-growing bacteria and several days for slower growing organisms[19–21]. Since reducing the time to optimal antimicrobial therapy significantly improves patient outcomes[22–24], rapid sequencing-based approaches that predict MICs may have clinical utility. The extensive WGS datasets generated by health agencies and the scientific community, such as nontyphoidal Salmonella strains, provides the necessary training sets required for building predictive models.

Several investigations have recently built models for predicting AMR phenotypes from WGS data. To date, the most common approach has relied on using a curated reference set of genes and polymorphisms that are implicated in AMR[25–33]. This reference-guided approach best predicts susceptibility and resistance when organisms are well studied and the AMR mechanisms are known. As larger collections of genomes have become available, several studies have used machine learning algorithms to predict susceptible and resistant phenotypes[27, 29, 31, 34–38]. By using WGS and AMR phenotype data to train a machine learning model, predictions without *a priori* information about the underlying gene content of the genome or molecular mechanism for resistance to each agent are possible. Although this reference-free approach requires many genomes, it is unbiased and can potentially be used to enable the discovery of new genomic features that are involved in AMR[36, 37]. These two general approaches have also been used to predict MICs for *Streptococcus, Neisseria*, and *Klebsiella*[35, 38–40]. When a curated reference collection of genes and SNPs is used for predicting MICs, a rules-based or machine learning algorithm is required for determining how much a given feature contributes to the MIC. Thus, for MIC prediction, both reference-guided and reference-free approaches are expected to have similar advantages and disadvantages if the collection of genes and SNPs used by the reference-guided method is sufficient for predicting all MICs, including those that are in the susceptible range. For example, in previous work, we built a machine learning model to predict MICs for a comprehensive population-based collection of 1,668 *Klebsiella pneumoniae* clinical isolates[38]. For each genome, we used nucleotide 10-mers and the MICs for each antibiotic as features to train the model. Extreme gradient boosting (XGBoost) was chosen as the machine learning algorithm[41]. The model could rapidly predict the MICs for 20 antibiotics with an average accuracy of 92%. This demonstrated that it is possible to successfully predict MICs without using a precompiled set of AMR genes or polymorphisms.

In this study, we build models that use whole genome sequence data to predict MICs for nontyphoidal *Salmonella* based on strains collected and sequenced by NARMS from 2002-2016. Our strategy can be used to guide responses to outbreaks and inform antibiotic stewardship decisions.

## Materials and Methods

### Genomes and Metadata

A total of 5,278 nontyphoidal *Salmonella* genome sequences were used in this study. All strains were collected and sequenced as part of the NARMS program. The strains were recovered from either raw retail meat and poultry or directly from livestock animals at slaughter. Antimicrobial susceptibility testing was performed using broth microdilution on the Sensititre^®^ system (Thermo Scientific) for 15 antibiotics: ampicillin (AMP), amoxicillin/clavulanic acid (AUG), ceftriaxone (AXO), azithromycin (AZI), chloramphenicol (CHL), ciprofloxacin (CIP), trimethoprim/sulfamethoxazole (COT), sulfisoxazole (FIS), cefoxitin (FOX), gentamicin (GEN), kanamycin (KAN), nalidixic acid (NAL), streptomycin (STR), tetracycline (TET), and ceftiofur (TIO) at FDA and USDA NARMS laboratories[13]. Clinical breakpoints are based on CLSI and FDA guidelines[42]. Whole genome sequencing was performed using the lllumina HiSeq and MiSeq platforms using standard methods[25]. Accession numbers and MICs for each isolate are listed in Table S1. All non-AMR metadata including serotypes, host, geographic location of isolation and isolation year were taken from the metadata associated with each NCBI SRA entry.

### Genomic Analyses

The short read sequence data for each strain was assembled with the PATRIC genome assembly service[43], using the “Full SPAdes” pipeline which uses BayesHammer[44] for read correction and SPAdes for assembly[45]. All genomes were annotated using the PATRIC annotation service[43], which uses a variation of the RAST tool kit annotation pipeline[46]. Annotated genomes are available on the PATRIC website (https://patricbrc.org). PATRIC genome identifiers are displayed in Table S1. Protein annotations, including those specifically asserted to be involved in AMR[47] were downloaded from the PATRIC workspace and used for subsequent analyses. A phylogenetic tree was generated for the strains in the analysis by aligning the genes for the beta and beta prime subunits of the RNA polymerase using MAFFT[48], concatenating the alignments, and computing a tree with FastTree[49]. The tree was rendered using iTOL[50].

### MIC Prediction

#### Model Generation

A model for predicting minimum inhibitory concentrations for the 15 antibiotics was built following the methods previously described by Nguyen and colleagues[38]. Briefly, each genome was divided into the set of nonredundant overlapping nucleotide 10-mers using the k-mer counting program KMC[51]. A matrix was built where the k-mers, antibiotics, and MICs are treated as features for each genome. Each row in the matrix contains the k-mers for a genome as well as the MIC for a single antibiotic. The MIC prediction model was built using an XGBoost[41] regressor predicting linearized MICs. All model parameters were identical to those used by Nguyen et al[38]. Ten-fold cross validations were used to assess the overall accuracy and sensitivity of every model used in this study. A non-overlapping training set (80% of the data), validation set (10% of the data), and test set (10% of the data) were generated for each fold. The validation set was used to monitor each model to prevent overfitting. Unless otherwise stated, the accuracy is reported as the ability to predict the correct MIC within ± 1 two-fold dilution step of the laboratory-derived MIC. Defining an accuracy to be within one two-fold dilution step is consistent with FDA requirements for automated MIC measuring device standards and is consistent with established clinical microbiology practices[20, 52, 53]. A comparison of raw accuracies and accuracies within ±1 two-fold dilution step is shown in Table S2. To assess the accuracy of a model over various metadata categories including date, serotype source, and location, the training set genomes are used to make the model. The test set genomes are used to assess the model accuracy for a given fold. For models based on date ranges, all parameters are identical and the accuracy is reported over the genomes from the held-out dates.

#### Subsampling

In order to perform the model building on a machine with 1.5 TB of RAM (machines with more memory are currently somewhat uncommon), we reduced the matrix size to sets of size *n*, where *n* ≤ 250, 500,1000, 2000, 3000, 4000, and 4500 genomes respectively. To create a diverse subset of size *n*, a hierarchical clustering method[54] was used to create *n* clusters by using the 10-mer distribution of each genome as input features. To avoid the curse of dimensionality[55, 56], the taxicab/Manhattan distance (*L*_1_ norm) was used, rather than the Euclidean distance (*L*_2_ norm), since previous research has shown it to be both computationally fast and more accurate for high dimensional data[57]. From the resulting *n* clusters, one genome from each cluster was randomly selected from a uniform distribution to create the subset containing *n* genomes. For each subset of genomes, a matrix was generated, and models were generated as described above.

#### Feature identification

In order to unambiguously identify k-mers that are important to MIC prediction, we built separate models for each individual antibiotic using the method described above, except that we increased the k-mer length to 15 nucleotides in order to reduce the number of redundant k-mers within each genome and to enable analyses with BLAST[58]. We also measured k-mer hits as presence versus absence, rather than counts, in order to simplify the analysis. Each model was built using the set of 1,000 diverse genomes from the subsampling experiment described above and 10-fold cross validations were performed on each model.

XGBoos’s internal feature importance was computed for each fold within the 10-fold cross validation. This results in an importance score per feature (15-mer) from each fold. In order to generate an overall importance score for the top features, we summed the feature importance scores from each fold for the top ten features. This overall importance score captures both the importance of the 15-mer to a given fold and the number of times that 15-mer was chosen as a top feature within each of the ten folds.

XGBoos’s internal feature importance is unable to provide correlations between features and label values, and thus does not provide an indication of whether a k-mer is related to antibiotic resistance or susceptibility. This is partially due to the fact that many non-linear correlations exist that may use multiple features. In order to see if the high scoring k-mers correlate with resistance or susceptibility, we computed the distribution of MICs for the genomes containing each high scoring k-mer. For example, a k-mer conferring susceptibility should be found in more genomes with lower MICs, while a k-mer conferring resistance should exist in genomes with higher MICs. Each high scoring k-mer was also compared to the set of protein encoding genes within each *Salmonella* genome. If a k-mer was found within a known AMR gene, that gene was reported. Otherwise, all protein-encoding genes within 3kb of the k-mer were reported in order to assess the neighborhood of the k-mer.

To find k-mers that are being used by the individual antibiotic models to predict susceptible MICs, we computed the presence or absence of each k-mers with high XGBoost feature importance values (described above) for the entire data set of 5278 genomes. The k-mers with the largest difference in occurrence between the susceptible and resistant genomes are the ones that are being chosen by the models for predicting susceptible MICs. To demine if there were significant SNPs in these k-mers, we found the genomic features containing the k-mer—protein encoding gene, RNA gene, or intergenic region—using BLASTn[58]. The corresponding feature or region was then found for all genomes in the collection. The features were aligned using MAFFT[48] and manually curated using Jalview[59]. Poor quality sequence was removed, all duplicates and paralogs were removed, and the subalignment covering the k-mer was extracted. To prevent possible biases due to clonality that may exist in the full set of genomes, the analysis was repeated on the diverse subset of 1000 genomes (described above). We report a SNP in a k-mer region as being significant if the susceptible and resistant sets are significantly different (P-value < 0.001) for a given nucleotide position based on a Chi-square test for both the full set of 5278 genomes and the set of 1000 diverse genomes. Sequence logos for k-mers containing significant SNPs were generated using WebLogo[60]. K-mers from the Azithromycin and Ciprofloxacin models were excluded from this analysis because they each had seven resistant genomes. Comparisons of codon usage were computed versus the genome average, genome mode, and high expression gene sets as described previously[61, 62].

### Software availability

The *Salmonella* MIC prediction model based on 4,500 genomes—including the software and documentation for running the model—is available at our GitHub page, https://github.com/PATRIC3/mic_prediction.

## Results

### Model Construction

For this study, we used a publicly available collection of 5,278 *Salmonella* whole genome sequences generated by the NARMS project between 2002 and 2016. The strains were isolated from retail meat and food animal sources in the United States. The collection includes 98 different serotypes, including Heidelberg (678 genomes), Kentucky (618 genomes), and Typhimurium var. 5- (588 genomes) from 41 states (Table S1). Isolates were tested for resistance to up to 15 antimicrobial agents using the broth microdilution method. Many of the strains were randomly selected for sequencing as part of a compulsory nation-wide collection program (Table 1).

**Table 1.** The number of susceptible, intermediate and resistant genomes across the 15 antibiotics for the 5278 *Salmonella* genomes used in this study.

| Antibiotic | Susceptible genomes | Intermediate genomes | Resistant genomes |
| --- | --- | --- | --- |
| AMP | 3682 | 2 | 1593 |
| AUG | 4145 | 355 | 778 |
| AXO | 4508 | 1 | 769 |
| AZI | 2409 | 0 | 7 |
| CHL | 5026 | 87 | 164 |
| CIP | 5217 | 53 | 7 |
| COT | 5219 | 0 | 58 |
| FIS | 3356 | 0 | 1573 |
| FOX | 4501 | 98 | 679 |
| GEN | 4577 | 68 | 633 |
| KAN | 837 | 3 | 84 |
| NAL | 5233 | 0 | 45 |
| STR | 872 | 0 | 1919 |
| TET | 2364 | 28 | 2885 |
| TIO | 4517 | 8 | 753 |

The nontyphoidal *Salmonella* MIC prediction model was built similar to our previously described strategy used to predict MICs for *K. pneumoniae* clinical isolates[38]. Since the *Salmonella* data set has many more genomes and greater sampling in the range of susceptible MICs, it provides a critical test case for determining if the approach remains robust for other pathogens. In the *Klebsiella* study, we built individual models for each antibiotic, as well as a single large integrated model by combining the data from all antibiotics. We found that the combined model achieved slightly higher overall accuracies (by ~l-2%), however the matrix that was necessary to train this model had a large memory footprint. Indeed, if we were to build a similar matrix for the current *Salmonella* data set using all 5,278 genomes, the model training would exceed 1.5 TB of RAM. Therefore, we first built models for all antibiotics using subsets of the genomes ranging in size from 250-4,500 genomes that were rationally selected to maximize genetic diversity (Figure 1). A matrix built from 4,500 genomes is the largest we can train on a 1.5 TB machine using this protocol. As the training set size increases from 250 to 1000 genomes, the accuracy increases from 88.5% to 91.4%. Then as the training set increases beyond 1000 genomes, the accuracy continues to improve more modestly, with the 4,500-genome model having an average accuracy of 95.2%. Results indicate that the overall MIC prediction approach, which was developed previously for *Klebsiella pneumoniae*, also works for *Salmonella* despite the differences in sampling, genetic diversity and MICs. Also, we discovered that a smaller number of well-chosen diverse genomes can serve as a useful proxy for representing the entire set, since models built from ≥500 genomes have accuracies exceeding 90%.

**Figure 1.**
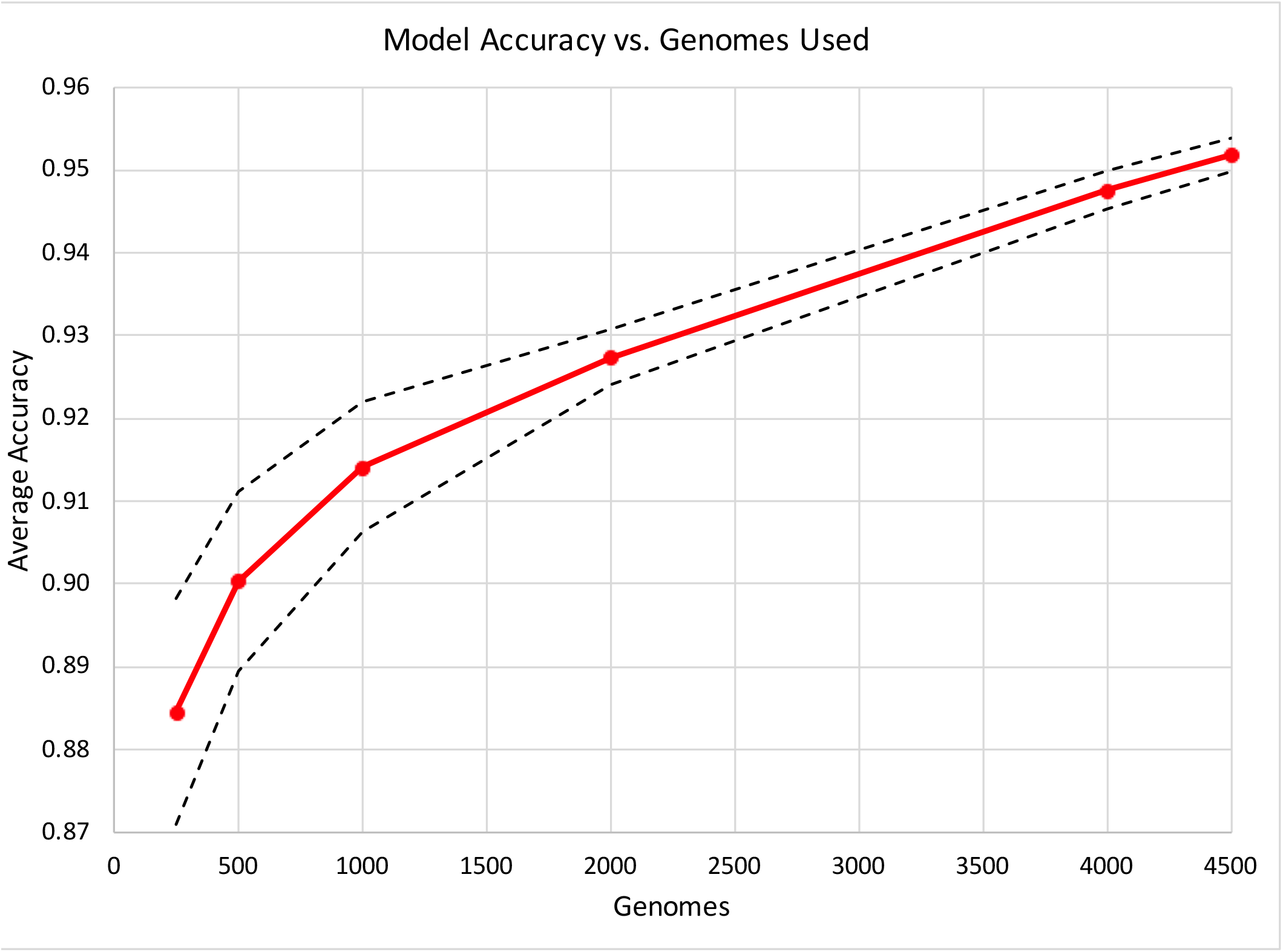
MIC prediction model accuracy for subsamples of genomes. Diverse subsamples of genomes were chosen and the model accuracy within ± 1 two-fold dilution step based on a 10-fold cross validation is shown with the red plot line. The dashed line represents the high and low values for the 95% confidence interval for the average accuracy at each given plot point.

### Model Accuracy

We computed the overall accuracy for each antibiotic using the model that is based on 4,500 genomes. For this model, all 15 antibiotics have average accuracies ≥90%, with their Q_1_ quartile bound ≥89% (Figure 2). Chloramphenicol and ceftiofur had the highest accuracies (99%), and gentamicin and tetracycline had the lowest accuracies (91% and 90%, respectively) (Table S2). Since the model is robust to the various mechanisms of resistance for the 15 antibiotics, it is possible that the slightly lower accuracies for gentamicin and tetracycline could be due to the distribution of multiple AMR genes/mechanisms across the population of strains with resistant genomes (which will be analyzed in more detail below). Figure 3 depicts the accuracy of the 4,500-genome model for each MIC. Overall, the model is robust for both the resistant and susceptible MICs, and it tends to be more accurate when a MIC is represented by many genomes. The model tends to have lower accuracies for the highest and lowest MICs, perhaps because of underlying genetic differences between strains that have been reported with ≥ or ≤ values, which represents a range of MICs rather than a discrete value.

**Figure 2.**
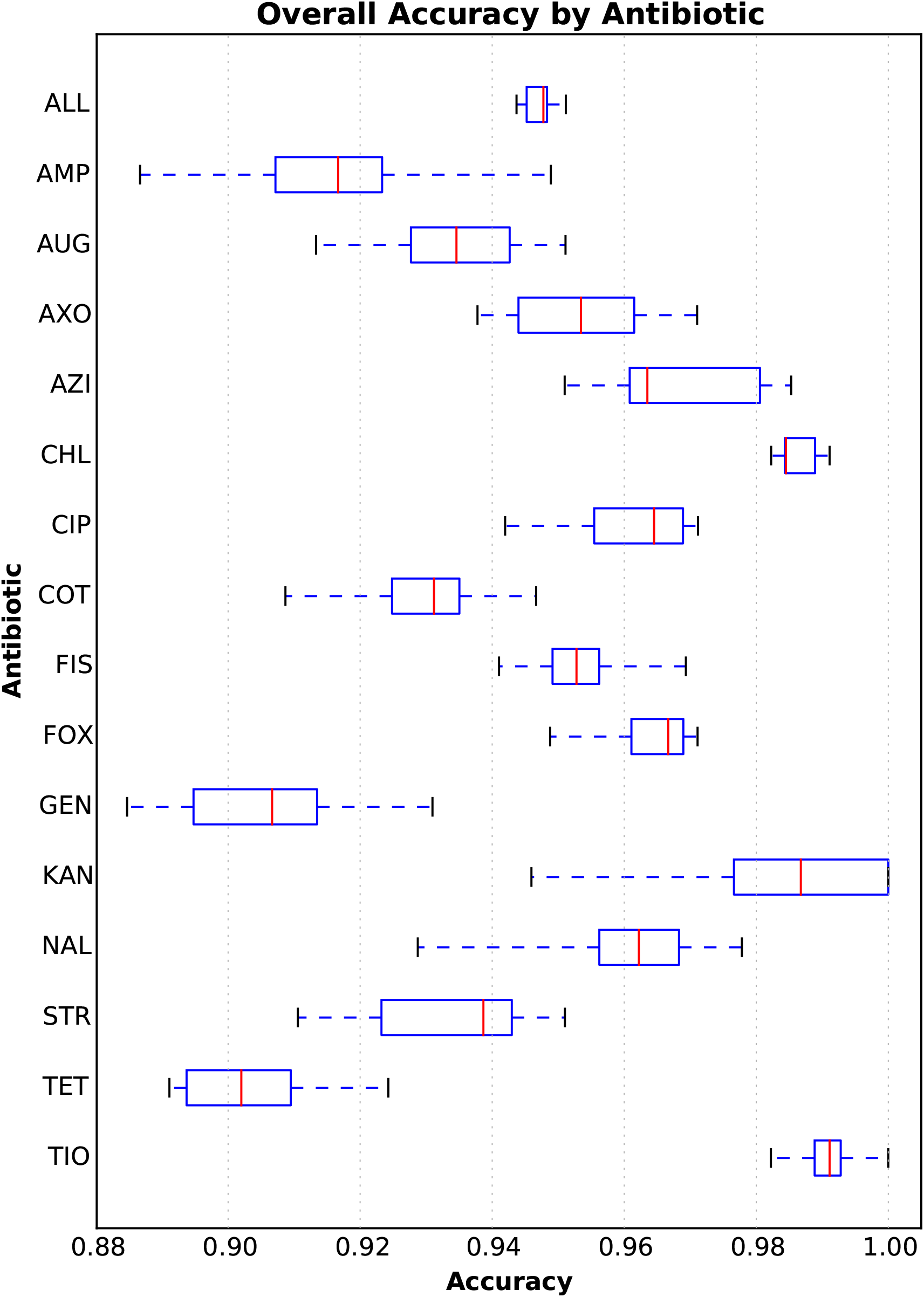
Box plot of the overall accuracies within ±1 two-fold dilution step for each antibiotic in the 4500-genome model. The Y-axis depicts each antibiotic (abbreviations are defined in Materials and Methods). The X-axis depicts the accuracy. Each vertical red line represents the median accuracy over the holdout sets for each fold in the ten-fold cross validation. The blue box encompasses the data of the first and third quartiles. The dashed blue horizontal lines bounded by black vertical lines (or “whiskers”) depict the entire distribution of accuracies for each fold and antibiotic. The accuracy of the entire 4500 genome model over all antibiotics and folds is depicted in the row marked “ALL”.

**Figure 3.**
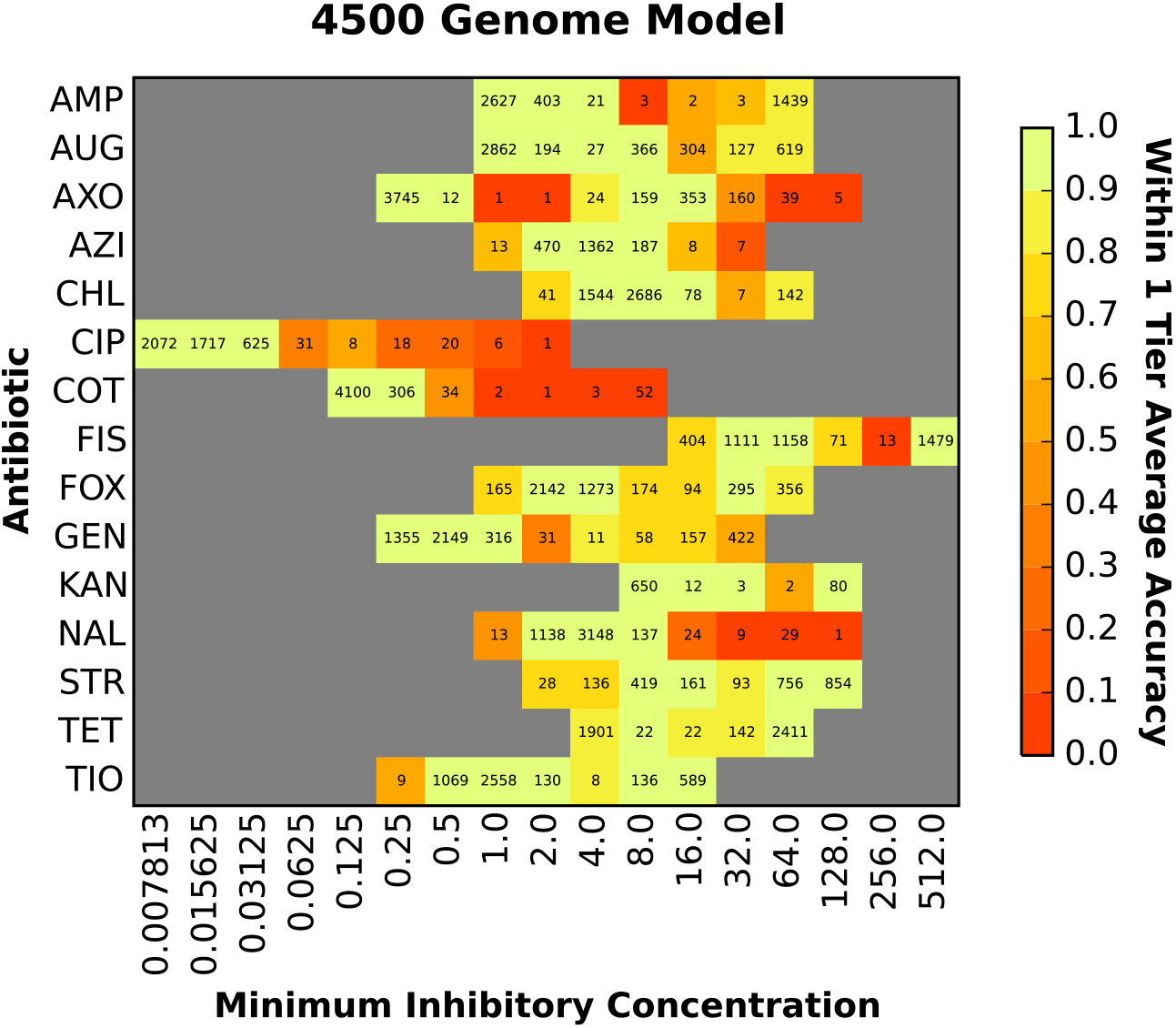
The accuracy of the MIC prediction model based on 4,500 diverse genomes. The heat map depicts the accuracy within ±1 two-fold dilution step of the laboratory-derived MIC. The X-axis shows the MIC (μg/ml) and each antibiotic is shown on the Y-axis. The accuracy for each antibiotic-MIC combination is depicted by color with bright yellow/green being the most accurate and red being the least accurate. The values shown in each cell are the number of genomes with that MIC for a given antibiotic.

The utility of AMR diagnostic devices is often described in terms of error rate. Major errors (MEs) are defined as susceptible genomes that have been incorrectly assigned resistant MICs by the model. Very major errors (VMEs) are defined as resistant genomes that have been incorrectly assigned susceptible MICs by the model. FDA standards for automated systems recommend a major error rate ≤ 3%[53]. All antibiotics used in the model have ME rates within this range (Table 2). The FDA standards for VME rates indicate that the lower 95% confidence limit should be ≤1.5% and upper limit should be ≤7.5%[53]. Seven of the 15 antibiotics—amoxicillin/clavulanic acid, ceftriaxone, chloramphenicol, cefoxitin, streptomycin, tetracycline and ceftiofur—have acceptable VME rates by this measure. Ampicillin and sulfisoxazole have VME rates with 95% confidence intervals approaching this range: [0.022, 0.033] and [0.026, 0.053] respectively. The VME rates are higher for some of the remaining antibiotics because there are fewer resistant genomes. As more resistant genomes are collected, and the data set becomes more balanced, we expect VME rates to be reduced.

**Table 2.** Very major error (VME) rate, defined as resistant genomes predicted as being susceptible, and major error (ME) rate, defined as susceptible genomes predicted as being resistant, for the 4500-genome model.

| Antibiotic | VME Avg<sup>1</sup> | VME 95% CI<sup>2</sup> | ME Avg<sup>1</sup> | ME 95% CI<sup>2</sup> | Resistant Samples | Susceptible Samples |
| --- | --- | --- | --- | --- | --- | --- |
| All | 0.027 | [0.024-0.030] | 0.001 | [0.001-0.002] | 10979 | 47366 |
| AMP | 0.028 | [0.022-0.033] | 0.000 | [0.000-0.001] | 1442 | 3054 |
| AUG | 0.012 | [0.000-0.025] | 0.000 | [0.000-0.000] | 746 | 3449 |
| AXO | 0.022 | [0.011-0.032] | 0.000 | [0.000-0.001] | 740 | 3758 |
| AZI | 0.857 | [0.508-1.207] | 0.000 | [0.000-0.000] | 7 | 2040 |
| CHL | 0.000 | [0.000-0.000] | 0.000 | [0.000-0.001] | 149 | 4271 |
| CIP | 0.417 | [-0.099-0.933] | 0.000 | [0.000-0.000] | 7 | 4445 |
| COT | 0.670 | [0.515-0.825] | 0.000 | [0.000-0.001] | 55 | 4443 |
| FIS | 0.039 | [0.026-0.053] | 0.000 | [0.000-0.000] | 1479 | 2757 |
| FOX | 0.009 | [-0.001-0.020] | 0.000 | [0.000-0.000] | 651 | 3754 |
| GEN | 0.090 | [0.066-0.113] | 0.000 | [0.000-0.000] | 579 | 3862 |
| KAN | 0.074 | [0.012-0.136] | 0.000 | [0.000-0.000] | 82 | 662 |
| NAL | 0.917 | [0.819-1.014] | 0.000 | [0.000-0.001] | 39 | 4460 |
| STR | 0.014 | [0.008-0.020] | 0.027 | [0.013-0.040] | 1703 | 744 |
| TET | 0.000 | [0.000-0.000] | 0.018 | [0.012-0.025] | 2575 | 1901 |
| TIO | 0.004 | [-0.001-0.009] | 0.000 | [0.000-0.000] | 725 | 3766 |

<sup>1</sup> Reported within $\pm 1$ two-fold dilution step
<sup>2</sup> 95% confidence interval

In addition to the extensive MIC data, NARMS reports rich metadata including isolation date, food or animal source, collection year, geographic location and serotype. We computed the accuracy of the model over each available metadata category to determine if the model is robust to these differences and to ensure that no subset is biasing the model. The genomes span a 15-year collection period, with all the years except 2002 (the oldest) and 2016 (the most recent) having over 100 isolates. The model accuracy ranges from 94-97% over each collection year (Table 3). That is, the genetic factors that contribute to the MICs have either remained stable over the 15-year period or have been learned as the model was trained. Although the data set is mostly comprised of poultry meat or live animal isolates, the accuracy ranges between 94-96% over the four contamination sources: turkey, beef, pork, and chicken (Table 4). No obvious biases were detected in the accuracies based on the state of isolation (an average of 95% accuracy over 41 states with a 95% Cl equal to [0.95-0.96]) (Figure 4) or the serovars of each isolate (94% accuracy over 97 serovars with a 95% Cl equal to [0.94-0.96]) (Table S3). Since the traditional *Salmonella* serotyping scheme is based the lipopolysaccharide O and flagellar H antigens, which are encoded by genes that influence the cell surface[63], we also constructed a phylogenetic tree for *Salmonella* genomes to observe the model accuracy over the various clades. Overall, no phylogenetic bias in the model accuracy was detected (Figure S1).

**Table 3.** Model accuracy for the genomes from each sample collection year.

| Collection Date | Accuracy | Genomes | Bins<sup>*</sup> |
| --- | --- | --- | --- |
| 2002 | 0.97 | 55 | 624 |
| 2003 | 0.95 | 159 | 1809 |
| 2004 | 0.96 | 235 | 2850 |
| 2005 | 0.95 | 274 | 3384 |
| 2006 | 0.95 | 313 | 3880 |
| 2007 | 0.94 | 258 | 3192 |
| 2008 | 0.95 | 388 | 4821 |
| 2009 | 0.95 | 436 | 5367 |
| 2010 | 0.94 | 230 | 2820 |
| 2011 | 0.95 | 214 | 2968 |
| 2012 | 0.96 | 257 | 3694 |
| 2013 | 0.97 | 265 | 3793 |
| 2014 | 0.95 | 506 | 7100 |
| 2015 | 0.95 | 689 | 9646 |
| 2016 | 0.96 | 83 | 1161 |

<sup>\*</sup>The total number of MICs available for the genomes isolated in that year

**Table 4.** Model accuracy for the genomes isolated from various sources.

| Source | Accuracy | Genomes | Bins<sup>*</sup> |
| --- | --- | --- | --- |
| Chicken | 0.96 | 1981 | 25869 |
| Cow/Beef | 0.94 | 419 | 5688 |
| Pig/Pork | 0.95 | 448 | 6144 |
| Turkey | 0.94 | 1651 | 21260 |

<sup>\*</sup>The total number of MICs available for the genomes of each category

**Figure 4.**
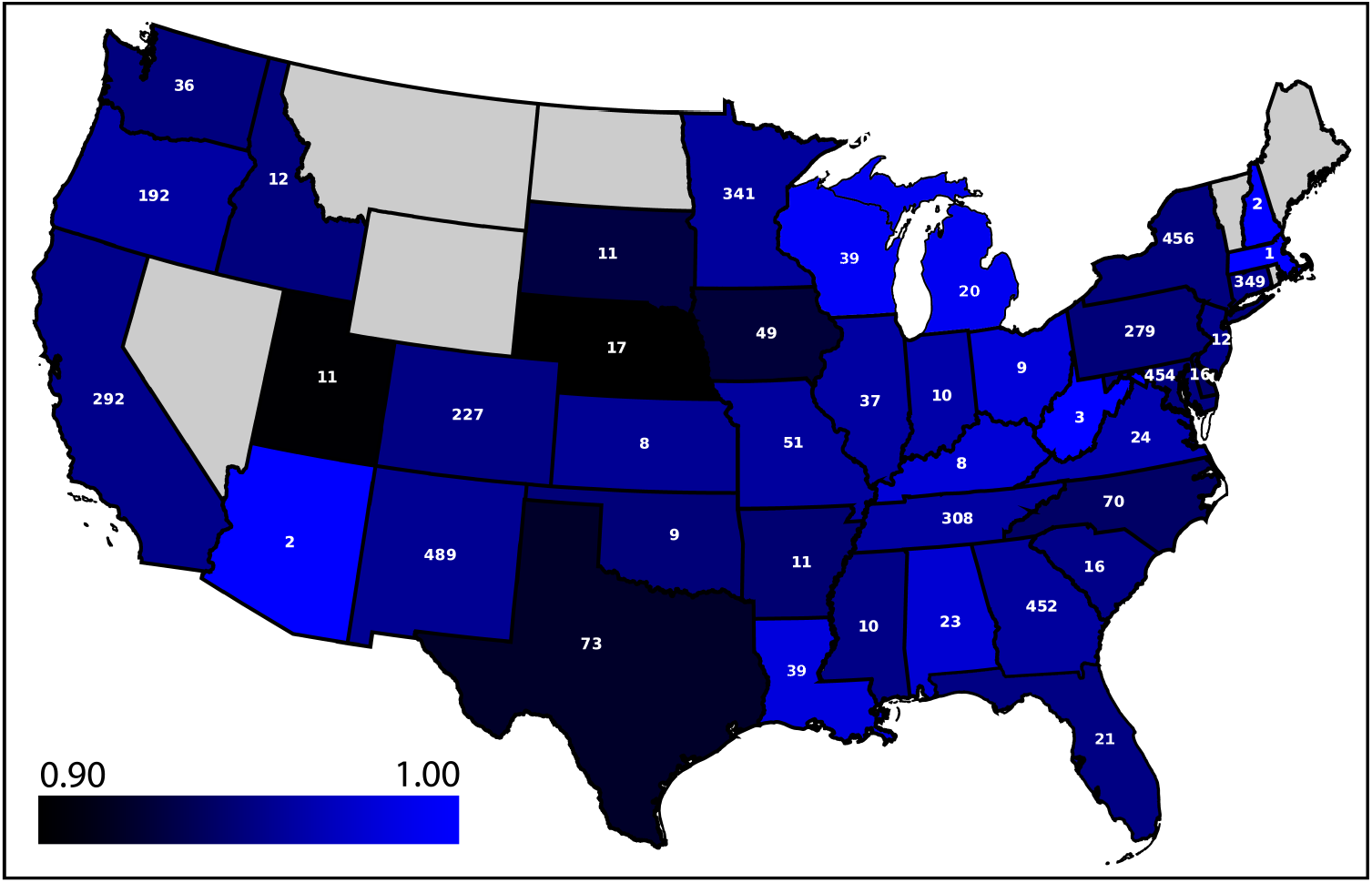
The average accuracy of the model based on 4,500 diverse genomes for predicting MICs for the *Salmonella* genomes from each state. Light blue is most accurate and dark blue/black is least accurate. Note that the scale starts at an accuracy of 0.90. Each state is labeled with the number of genomes collected from that state. States without a label contain no samples and are colored in grey; no genomes exist in the collection from Alaska and Hawaii.

One concern of using a model that is trained on the data from previous years, in some cases over 15 years old, is that the training set is not representative of currently circulating strains. That is, the model may be inaccurate for predicting MICs for genomes of strains that are currently circulating or will emerge in the future. For example, shifts in clonal groups, evolution of AMR-associated genes, or introduction of AMR genes by horizontal gene transfer is possible[64, 65]. We evaluated this possibility by building models from subsets of the whole genome sequence data using strains collected in earlier years and measuring the accuracy of the models on genomes collected in later years. Models were built for years prior to 2009 through 2014 and tested on the remaining genomes (Table 5). These models have accuracies ranging from 86-92%. As the number of years used for building the models increases, the number of genomes available for testing decreases, so we also tested each model on only the 462 genomes from 2015 and 2016. Similarly, the accuracy of each model on the 2015 and 2016 genomes ranges from 87-90% (Table S4). The results indicate that within this data set, models generated from genomes collected at earlier dates yield stable MIC predictions for genomes collected at later dates. This finding is consistent with the pattern of AMR genes that is observed within the data set. Although AMR gene content may vary from year to year, we do not observe any major sweeps or fixation events that drastically alter the AMR gene content of the collection between years, which would cause the MIC predictions to fail for a large fraction of the genomes (Table S5). Taken together, these data suggest that the MIC prediction models generated in this study are likely to be sustainable over time.

**Table 5.** The ability of models trained on genomes from prior years to predict MICs for genomes collected in later years.

| **Training set years** | **Test set years** | **Accuracy** | **95% CI** | **Training Bins \*** | **Testing Bins \*** | **Training Genomes** | **Testing Genomes** |
| --- | --- | --- | --- | --- | --- | --- | --- |
| 2002-2008 | 2009-2016 | 0.88 | [0.88-0.89] | 36563 | 22412 | 1819 | 2681 |
| 2002-2009 | 2010-2016 | 0.88 | [0.88-0.89] | 31196 | 27779 | 2255 | 2245 |
| 2002-2010 | 2011-2016 | 0.88 | [0.88-0.88] | 28376 | 30599 | 2485 | 2015 |
| 2002-2011 | 2012-2016 | 0.88 | [0.88-0.89] | 25408 | 33567 | 2699 | 1801 |
| 2002-2012 | 2013-2016 | 0.88 | [0.87-0.88] | 21714 | 37261 | 2956 | 1544 |
| 2002-2013 | 2014-2016 | 0.86 | [0.86-0.87] | 17921 | 41054 | 3221 | 1279 |
| 2002-2014 | 2015-2016 | 0.92 | [0.92-0.92] | 10807 | 48168 | 3728 | 772 |

\*The total number of genome/antibiotic combinations

### Genomic regions important for MIC prediction

The 4,500-genome model described above contains data from all antibiotics and MICs, making feature extraction to determine which k-mers contribute to the MIC predictions for each antibiotic difficult. To address this limitation, we modified the protocol by building separate models for each antibiotic. Instead of using 10-mers, we increased the k-mer length to 15 nucleotides to reduce redundancy and make them identifiable using BLAST[58]. We also searched for presence or absence of k-mers, rather than using k-mer counts, to simplify the analysis of the XGBoost decision trees. Since a 15-mer matrix can be 4^5^ times larger than a 10-mer matrix, we used <= 1000 diverse genomes to reduce the memory footprint during training. Overall, the average accuracy for the individual models is nearly identical to the average accuracy for the combined 4,500-genome model (96% vs. 95%, respectively), and in nearly all cases, the 95% confidence intervals overlap between the combined and single antibiotic models (Table S6). Thus, for this data set, single antibiotic models with fewer genomes and larger k-mers perform as well as a combined model (Figure S2).

During model training, XGBoost assigns an importance value to each k-mer used in a decision tree. When the model is used to predict the MICs for a new genome, the k-mers with the highest importance values are the most informative for the MIC prediction. Thus, by analyzing the feature importance values of each k-mer, we can use the models as a tool for understanding the genomic regions that differentiate MICs. For each antibiotic-specific model, we parsed the XGBoost decision trees from each fold of the ten-fold cross validation to extract the importance values for each k-mer. To understand the relationship between known AMR genes and the important k-mers that were chosen by each model, we then searched for k-mers with high importance values within AMR genes that occur in close proximity to an AMR gene (in this case, we consider a window of 3kb, approximately 3 genes, to be a close proximity). Table 6 lists the highest-ranking k-mers from each model that occur within or in close proximity to an AMR gene. In most cases, the k-mers correspond to known AMR genes including class A and C beta-lactamases for the beta lactam antibiotics, aminoglycoside nucleotidyl- and acetyltransferases for the aminoglycosides, DNA gyrase and QnrB for the fluoroquinolones, TetA and TetR for tetracycline, and dihydrofolate reductase and dihydropteroate synthase for co-trimoxazole and sulfisoxazole. In the case of azithromycin, the collection contains mostly susceptible genomes (Table 1), so the first macrolide resistance gene observed corresponds with the eighth ranking k-mer. The top ten k-mers with the highest feature importance values from each of the ten folds used in model training are listed in Tables S7-S21. In addition to the top AMR k-mers displayed in Table 6, these tables show other highly ranking k-mers from the same AMR genes as well as k-mers from related genes that are known to confer resistance to the given antibiotic. In some cases, k-mers matching regions or genes from unrelated AMR mechanisms have high importance values, suggesting a pattern of co-occurrence on horizontally transferred genetic elements.

**Table 6.**
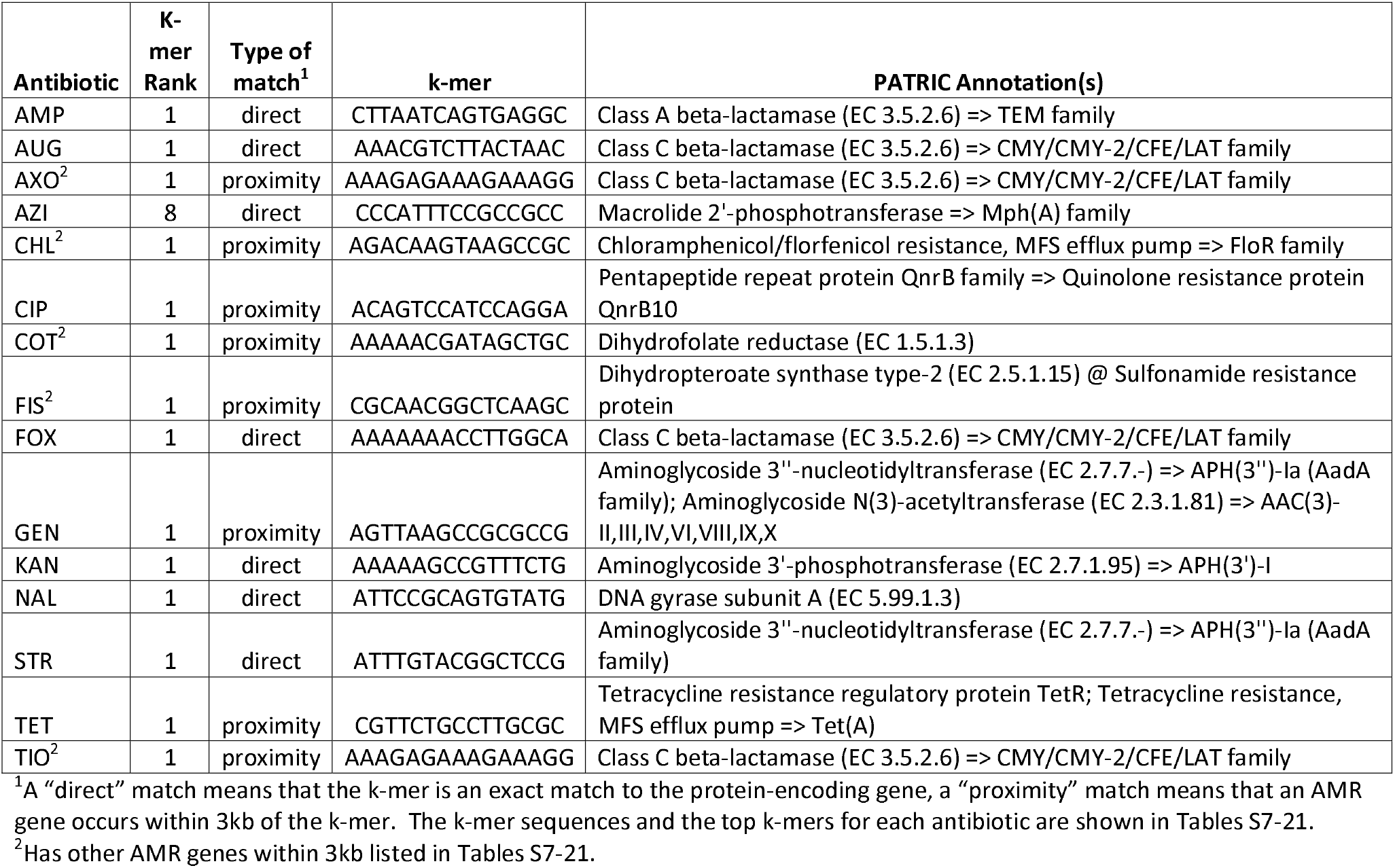
The highest-ranking AMR-related protein function (or genomic region) with a matching k-mer from the XGBoost models.

Since each model is predicting the entire range of MICs, some of the highly ranking k-mers will be used to predict susceptible MICs. To assess this, we computed the fraction of susceptible and resistant genomes with each k-mer from Tables S7-21. The set of k-mers that are most enriched in the susceptible genomes is shown in Table 7. Overall, seven of the top ten k-mers represent significantly different SNPs (P-value < 0.001) in both the complete set of 5,278 genomes and in the set of 1,000 diverse genomes used to build the models (Figure S3). The top k-mer associated with susceptibility is from the nalidixic acid model and occurs in the DNA gyrase *gyrA* gene. This is also the top k-mer that was found in an AMR gene for nalidixic acid from Table 6. In this case, the model is relying more heavily on the “wild type” version of the k-mer rather than any of the resistant versions (the remaining k-mers from Table 6 occur almost exclusively in resistant genomes). The same *gyrA* k-mer is also found as a highly ranking k-mer in the case of ciprofloxacin (Table S12). Two significant *gyrA* SNPs are captured by this k-mer (Figure S3). These are missense mutations in the resistant genomes occurring at Ser-83 and Asp-87, and changes at these positions have been shown to confer quinolone resistance in in *E. coli* [66, 67]. The remaining significant mutations from Figure S3 that occur in the protein-encoding genes are same sense (not amino acid changing) mutations. In the cases of *eptA* (Ser, TCG to TCA), *oadA* (Ala, GCC to GCA), the AraJ precursor gene (Leu, CTG to CTA), and the second *gcd* mutation (Thr, ACG to ACA), the codon changes from a commonly used codon in the susceptible genomes to the least preferred codon in the resistant genomes. In the cases of the *nrfE/nrfF* mutation (Asn, AAT to AAC) and the first *gcd* mutation (Asp, GAC to GAT), the resistant genomes have the preferred codon of the pair. Whether these SNPs have a modulating effect on protein translation or contribute to the fitness of the resistant organisms requires further analysis.

**Table 7.**
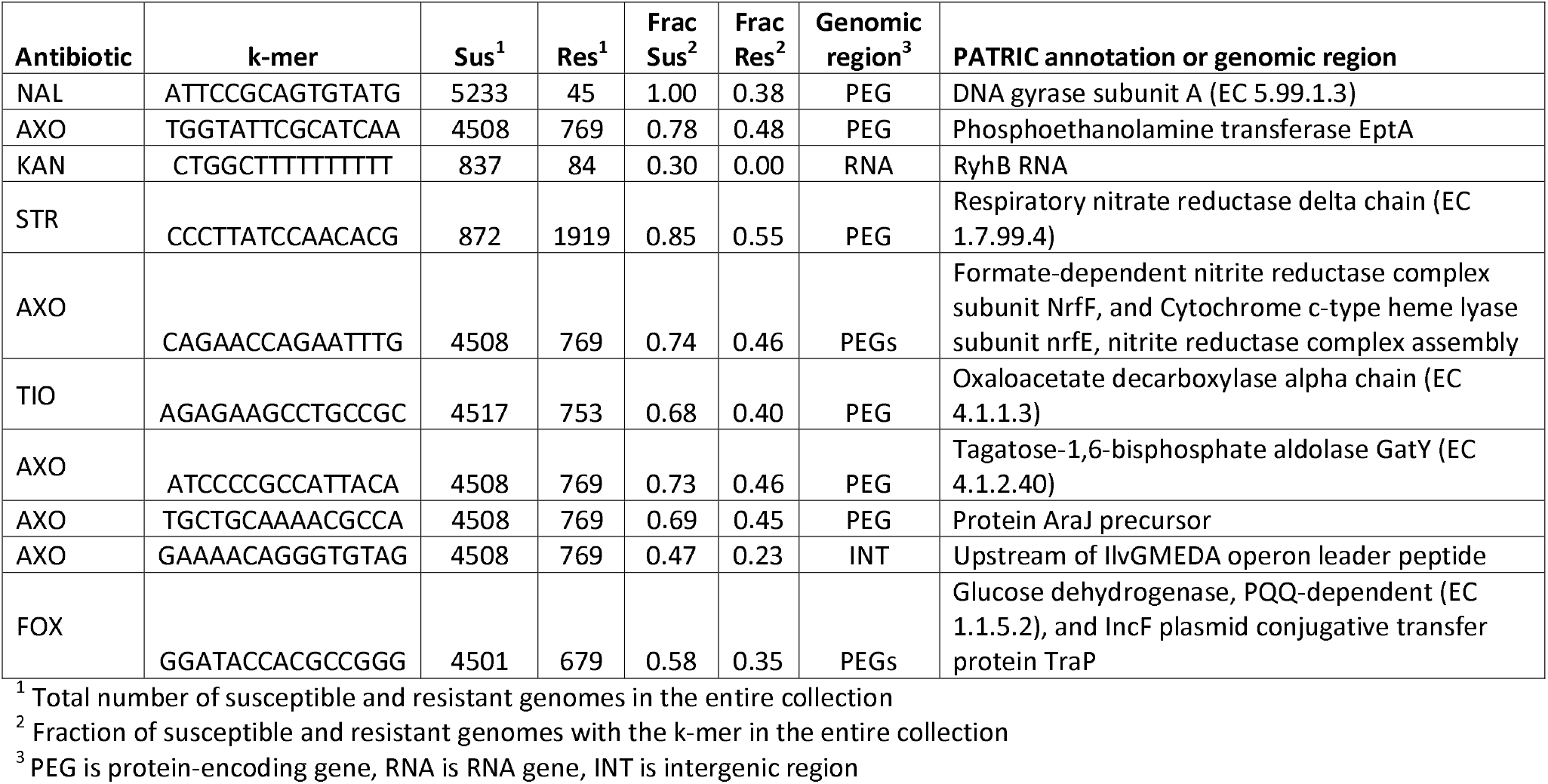
Important k-mers used by the individual antibiotic models for predicting susceptible MICs.

## Discussion

In this study, we have built machine learning-based MIC prediction models for nontyphoidal *Salmonella* genomes using XGBoost[41] that achieve overall accuracies of 95-96% within ± 1 two-fold dilution factor. To our knowledge, this is one of the largest and most accurate MIC prediction models to be published to date. Importantly, it provides a model strategy for performing MIC prediction directly from genome sequence data that could be applied to other human or veterinary pathogens.

The success of our MIC prediction model was dependent on the large, publicly available, population-based collection of genomes with associated metadata. Since researchers often focus on collecting highly resistant or otherwise unusual strains, the opportunities to generate balanced models are rare. We demonstrate the many benefits from comprehensive sampling for the entire range of possible MICs. First, diverse and balanced data sets improve model accuracies because there is better sampling across all MIC dilutions. Second, having balanced data enabled us to achieve acceptable ME and VME rates for 7 of the 15 antibiotics used in the study. Third, compared with our recent model for *Klebsiella pneumoniae*, the larger and more balanced data set in nontyphoidal *Salmonella* enabled us to build models for individual antibiotics that had similar accuracies to the combined model. This enabled us to begin to disambiguate the important genomic regions relating to resistant and susceptible MICs. Finally, we show that MICs in the susceptible range can be accurately predicted with the algorithm using all genomic data rather than scoping it to known AMR genes or gene polymorphisms. This contrasts with prior work correlating MICs to known resistance mechanisms in *Salmonella*[68]. In future studies, our strategy could be used as a starting point for identifying the subtle genomic changes that result in different MICs.

For each single-antibiotic model, we analyzed the k-mers that had high feature importance values and were important to the models for predicting MICs. The highly ranking k-mers that were enriched in the resistant genomes mainly occurred within or in close proximity to well-known AMR genes. With the exception of the *gyrA* k-mer, the highly ranking k-mers that were enriched in the susceptible genomes were significant in several cases, but more difficult to interpret. Some of these susceptibility k-mers hint at a possible relationship between AMR and oxidative stress or electron transport, such as the k-mers matching components of the nitrate and nitrite reductases and pqq-dependent glucose dehydrogenase, which is consistent with the known link between antibiotics to oxidative stress[69, 70]. Determining the molecular mechanisms underlying the susceptibility k-mers and AMR phenotypes should be further investigated.

The genomes in this study were collected over a 15-year period from 41 U.S. states. By building models encompassing ranges of earlier dates, we demonstrated stable and accurate MIC prediction for genomes collected at later dates. Presently, we are not aware of any large publicly available collections of *Salmonella* genomes with MIC data from other countries. Since AMR gene content may vary across pathogen populations, validation of the *Salmonella* models using strains from other countries is important to its application in global health. Nevertheless, the present analysis clearly demonstrates that current model provides accurate MIC predictions for United States isolates. Similarly, an analysis of this model on *Salmonella typhi* strains would provide information about the utility of the model over broader phylogenetic distances.

## Acknowledgements

This work was supported by the National Institute of Allergy and Infectious Diseases, National Institutes of Health, Department of Health and Human Service [Contract No. HHSN272201400027C]. We thank Emily Dietrich for her helpful edits.

## Author contribution statement

MN: study design, experiments, data generation manuscript preparation, SWL: study design, PFM: study design, data generation, RJO: study design, RO: software engineering, RLS: study design, GHT: study design, data generation, SZ: study design, data generation, JJD: study design, data generation, manuscript preparation

## Additional Information

### Accession Codes

Data are available under bioprojects PRJNA292661 and PRJNA292666. SRA run accession for each genome are displayed in (Table SI).

### Competing financial interests

The authors claim no competing financial interests.

## Disclaimer

The views expressed in this article are those of the authors and do not necessarily reflect the official policy of the Department of Health and Human Services, the U.S. Food and Drug Administration, and Centers for Disease Control and Prevention or the U.S. Government. Mention of trade names or commercial products in this publication is solely for the purpose of providing specific information and does not imply recommendation or endorsement by the U.S. Department of Agriculture or Food and Drug Administration.

## Abbreviations

AMP: ampicillin
AMR: antimicrobial resistance
AUG: amoxicillin/clavulanic acid (Augmentin)
AXO: ceftriaxone AZI: azithromycin
CDC: United States Centers for Disease Control and Prevention
CHL: chloramphenicol CIP: ciprofloxacin
CLSI: Clinical and Laboratory Standards Institute
COT: trimethoprim/sulfamethoxazole (co-trimoxazole)
FDA: United States Food and Drug Administration
FIS: sulfisoxazole
FOX: cefoxitin
FSIS: USDA Food Safety and Inspection Service
GEN: gentamicin
KAN: kanamycin
ME: major error
MIC: minimum inhibitory concentration
NAL: nalidixic acid
NARMS: National Antimicrobial Resistance Monitoring System
SNP: single nucleotide polymorphism
STR: streptomycin
TET: tetracycline
TIO: ceftiofur
USDA: United States Department of Agriculture
VME: very major error
WGS: whole genome sequencing
XGBoost: Extreme Gradient Boosting

## References

1. Centers for Disease Control and Prevention (CDC). Surveillance for Foodborne Disease Outbreaks, United States, 2015, Annual Report. Atlanta, Georgia: US Department of Health and Human Services, CDC. 2017. Available from: https://www.cdc.gov/foodsafetv/pdfs/2015FoodBomeQutbreaks508.pdf.

2. Crim SM, Griffin PM, Tauxe R, Marder EP, Gilliss D, Cronquist AB, et al. Preliminary incidence and trends of infection with pathogens transmitted commonly through food- Foodborne Diseases Active Surveillance Network, 10 US sites, 2006–2014. MMWR Morbidity and mortality weekly report. 2015;64(18):495–9.

3. Scallan E, Hoekstra RM, Widdowson M, Hall A, Griffin P. Foodborne illness acquired in the United States. Emerging Infectious Diseases. 2011;17(7):1339–40.

4. World Health Organization. WHO estimates of the global burden of foodborne diseases: foodborne disease burden epidemiology reference group 2007–2015. 2015.

5. Andino A, Hanning I. *Salmonella enterica*: survival, colonization, and virulence differences among serovars. The Scientific World Journal. 2015;2015.

6. Aserkoff B, Bennett JV. Effect of antibiotic therapy in acute salmonellosis on the fecal excretion of salmonellae. N Engl J Med. 1969;281(12):636–40. Epub 1969/09/18. doi: 10.1056/NEJM196909182811202.

7. Crump JA, Sjölund-Karlsson M, Gordon MA, Parry CM. Epidemiology, clinical presentation, laboratory diagnosis, antimicrobial resistance, and antimicrobial management of invasive *Salmonella* infections. Clinical microbiology reviews. 2015;28(4):901–37.

8. Varma JK, Mølbak K, Barrett TJ, Beebe JL, Jones TF, Rabatsky-Ehr T, et al. Antimicrobial-resistant nontyphoidal *Salmonella* is associated with excess bloodstream infections and hospitalizations. The Journal of infectious diseases. 2005;191(4):554–61.

9. Varma JK, Greene KD, Ovitt J, Barrett TJ, Medalla F, Angulo FJ. Hospitalization and antimicrobial resistance in *Salmonella* outbreaks, 1984–2002. Emerging Infectious Diseases. 2005;11(6):943.

10. Krueger AL, Greene SA, Barzilay EJ, Henao O, Vugia D, Hanna S, et al. Clinical outcomes of nalidixic acid, ceftriaxone, and multidrug-resistant nontyphoidal *Salmonella* infections compared with pansusceptible infections in FoodNet sites, 2006–2008. Foodborne pathogens and disease. 2014;11(5):335–41.

11. Angulo FJ, Mølbak K. Human health consequences of antimicrobial drug—resistant *Salmonella* and other foodborne pathogens. Clinical infectious diseases. 2005;41(11):1613–20.

12. Karp BE, Tate H, Plumblee JR, Dessai U, Whichard JM, Thacker EL, et al. National Antimicrobial Resistance Monitoring System: Two Decades of Advancing Public Health Through Integrated Surveillance of Antimicrobial Resistance. Foodborne pathogens and disease. 2017;14(10):545–57.

13. Food and Drug Administration (FDA). NARMS Now. Rockville, MD: 2018. Available from: https://www.fda.gov/AnimalVeterinary/SafetyHealth/AntimicrobialResistance/NationalAntimicrobialResistanceMonitoringSvstem/ucm416741.htm.

14. Abrams AJ, Trees DL. Genomic sequencing of *Neisseria gonorrhoeae* to respond to the urgent threat of antimicrobial-resistant gonorrhea. Pathogens and disease. 2017;75(4).

15. Goldberg B, Sichtig H, Geyer C, Ledeboer N, Weinstock GM. Making the leap from research laboratory to clinic: challenges and opportunities for next-generation sequencing in infectious disease diagnostics. MBio. 2015;6(6):e01888–15.

16. Didelot X, Bowden R, Wilson DJ, Peto TE, Crook DW. Transforming clinical microbiology with bacterial genome sequencing. Nature Reviews Genetics. 2012;13(9):601.

17. Brown EW, Gonzalez-Escalona N, Stones R, Timme R, Allard MW. The Rise of Genomics and the Promise of Whole Genome Sequencing for Understanding Microbial Foodborne Pathogens. Foodborne Pathogens: Springer; 2017. p. 333–51.

18. McArthur AG, Tsang KK. Antimicrobial resistance surveillance in the genomic age. Annals of the New York Academy of Sciences. 2017;1388(1):78–91.

19. Opota O, Croxatto A, Prod’hom G, Greub G. Blood culture-based diagnosis of bacteraemia: state of the art. Clinical Microbiology and Infection. 2015;21(4):313–22.

20. Reller LB, Weinstein M, Jorgensen JH, Ferraro MJ. Antimicrobial susceptibility testing: a review of general principles and contemporary practices. Clinical infectious diseases. 2009;49(11): 1749–55.

21. Saha SK, Darmstadt GL, Baqui AH, Hanif M, Ruhulamin M, Santosham M, et al. Rapid identification and antibiotic susceptibility testing of *Salmonella enterica* serovar Typhi isolated from blood: implications for therapy. Journal of clinical microbiology. 2001;39(10):3583–5.

22. Llor C, Bjerrum L. Antimicrobial resistance: risk associated with antibiotic overuse and initiatives to reduce the problem. Therapeutic advances in drug safety. 2014;5(6):229–41.

23. Kumar A, Roberts D, Wood KE, Light B, Parrillo JE, Sharma S, et al. Duration of hypotension before initiation of effective antimicrobial therapy is the critical determinant of survival in human septic shock. Critical care medicine. 2006;34(6):1589–96.

24. Palmer H, Palavecino E, Johnson J, Ohl C, Williamson J. Clinical and microbiological implications of time-to-positivity of blood cultures in patients with Gram-negative bacilli bacteremia. European journal of clinical microbiology & infectious diseases. 2013;32(7):955–9.

25. McDermott PF, Tyson GH, Kabera C, Chen Y, Li C, Folster JP, et al. Whole-genome sequencing for detecting antimicrobial resistance in nontyphoidal *Salmonella*. Antimicrobial agents and chemotherapy. 2016;60(9):5515–20.

26. Hunt M, Mather AE, Sánchez-Busó L, Page AJ, Parkhill J, Keane JA, et al. ARIBA: rapid antimicrobial resistance genotyping directly from sequencing reads. Microbial genomics. 2017;3(10).

27. Niehaus KE, Walker TM, Crook DW, Peto TE, Clifton DA, editors. Machine learning for the prediction of antibacterial susceptibility in *Mycobacterium tuberculosis*. 2014 IEEE-EMBS International Conference on Biomedical and Health Informatics (BHI); 2014: IEEE.

28. Stoesser N, Batty E, Eyre D, Morgan M, Wyllie D, Del Ojo Elias C, et al. Predicting antimicrobial susceptibilities for *Escherichia coli* and *Klebsiella pneumoniae* isolates using whole genomic sequence data. Journal of Antimicrobial Chemotherapy. 2013;68(10):2234–44.

29. Pesesky MW, Hussain T, Wallace M, Patel S, Andleeb S, Burnham C-AD, et al. Evaluation of machine learning and rules-based approaches for predicting antimicrobial resistance profiles in Gram-negative Bacilli from whole genome sequence data. Frontiers in microbiology. 2016;7:1887.

30. Lipworth SIW, Hough N, Leach L, Morgan M, Jeffrey K, Andersson M, et al. Whole genome sequencing for predicting *Mycobacterium abscessus* drug susceptibility. bioRxiv. 2018:251918.

31. Bradley P, Gordon NC, Walker TM, Dunn L, Heys S, Huang B, et al. Rapid antibiotic-resistance predictions from genome sequence data for *Staphylococcus aureus* and *Mycobacterium tuberculosis*. Nature communications. 2015;6:10063.

32. Harrison OB, Clemence M, Dillard JP, Tang CM, Trees D, Grad YH, et al. Genomic analyses of *Neisseria gonorrhoeae* reveal an association of the gonococcal genetic island with antimicrobial resistance. Journal of Infection. 2016;73(6):578–87.

33. Grad YH, Harris SR, Kirkcaldy RD, Green AG, Marks DS, Bentley SD, et al. Genomic epidemiology of gonococcal resistance to extended-spectrum cephalosporins, macrolides, and fluoroquinolones in the United States, 2000–2013. The Journal of infectious diseases. 2016;214(10):1579–87.

34. Coelho JR, Carriço JA, Knight D, Martinez J-L, Morrissey I, Oggioni MR, et al. The use of machine learning methodologies to analyse antibiotic and biocide susceptibility in *Staphylococcus aureus*. PLoS One. 2013;8(2):e55582.

35. Eyre DW, De Silva D, Cole K, Peters J, Cole MJ, Grad YH, et al. WGS to predict antibiotic MICs for *Neisseria gonorrhoeae*. Journal of Antimicrobial Chemotherapy. 2017;72(7):1937–47.

36. Drouin A, Giguère S, Déraspe M, Marchand M, Tyers M, Loo VG, et al. Predictive computational phenotyping and biomarker discovery using reference-free genome comparisons. BMC genomics. 2016;17(1):754.

37. Davis JJ, Boisvert S, Brettin T, Kenyon RW, Mao C, Olson R, et al. Antimicrobial resistance prediction in PATRIC and RAST. Scientific reports. 2016;6:27930.

38. Nguyen M, Brettin T, Long SW, Musser JM, Olsen RJ, Olson R, et al. Developing an in silico minimum inhibitory concentration panel test for *Klebsiella pneumoniae*. Scientific reports. 2018;8(1):421.

39. Metcalf BJ, Chochua S, Gertz R, Li Z, Walker H, Tran T, et al. Using whole genome sequencing to identify resistance determinants and predict antimicrobial resistance phenotypes for year 2015 invasive pneumococcal disease isolates recovered in the United States. Clinical Microbiology and Infection. 2016;22(12):1002. e1-. e8.

40. Li Y, Metcalf BJ, Chochua S, Li Z, Gertz RE, Walker H, et al. Penicillin-binding protein transpeptidase signatures for tracking and predicting β-lactam resistance levels in *Streptococcus pneumoniae*. MBio. 2016;7(3):e00756–16.

41. Chen T, Guestrin C, editors. XGBoost: A scalable tree boosting system. Proceedings of the 22nd acm sigkdd international conference on knowledge discovery and data mining; 2016: ACM.

42. US Food and Drug Administration (FDA). National Antimicrobial Resistance Monitoring System-Enteric Bacteria (NARMS): 2011 executive report. US Department of Health and Human Services. Food and Drug Administration, Rockville, MD. 2013.

43. Wattam AR, Davis JJ, Assaf R, Boisvert S, Brettin T, Bun C, et al. Improvements to PATRIC, the all-bacterial bioinformatics database and analysis resource center. Nucleic acids research. 2016;45(D1):D535–D42.

44. Nikolenko SI, Korobeynikov Al, Alekseyev MA, editors. BayesHammer: Bayesian clustering for error correction in single-cell sequencing. BMC genomics; 2013: BioMed Central.

45. Bankevich A, Nurk S, Antipov D, Gurevich AA, Dvorkin M, Kulikov AS, et al. SPAdes: a new genome assembly algorithm and its applications to single-cell sequencing. Journal of computational biology. 2012;19(5):455–77.

46. Brettin T, Davis JJ, Disz T, Edwards RA, Gerdes S, Olsen GJ, et al. RASTtk: a modular and extensible implementation of the RAST algorithm for building custom annotation pipelines and annotating batches of genomes. Scientific reports. 2015;5:8365.

47. Antonopoulos DA, Assaf R, Aziz RK, Brettin T, Bun C, Conrad N, et al. PATRIC as a unique resource for studying antimicrobial resistance. Briefings in bioinformatics. 2017.

48. Katoh K, Standley DM. MAFFT multiple sequence alignment software version 7: improvements in performance and usability. Molecular biology and evolution. 2013;30(4):772–80.

49. Price MN, Dehal PS, Arkin AP. FastTree 2-approximately maximum-likelihood trees for large alignments. PloS one. 2010;5(3):e9490.

50. Letunic I, Bork P. Interactive Tree Of Life (iTOL): an online tool for phylogenetic tree display and annotation. Bioinformatics. 2006;23(1):127–8.

51. Deorowicz S, Kokot M, Grabowski S, Debudaj-Grabysz A. KMC 2: fast and resource-frugal k-mer counting. Bioinformatics. 2015;31(10):1569–76.

52. Jorgensen JH. Selection criteria for an antimicrobial susceptibility testing system. Journal of clinical microbiology. 1993;31(11):2841.

53. US Food and Drug Administration (FDA). Class II Special Controls Guidance Document: Antimicrobial Susceptibility Test (AST) Systems. Rockville, MD: US FDA. 2009.

54. Pedregosa F, Varoquaux G, Gramfort A, Michel V, Thirion B, Grisel O, et al. Scikit-learn: Machine Learning in Python. Journal of Machine Learning Research. 2011;12:2825–30.

55. Bellman R. Dynamic programming. Princeton: Princeton University Press; 2013.

56. Shalev-Shwartz S, Ben-David S. Understanding machine learning: From theory to algorithms: Cambridge University Press; 2014.

57. Aggarwal CC, Hinneburg A, Keim DA, editors. On the surprising behavior of distance metrics in high dimensional space. International conference on database theory; 2001: Springer.

58. Camacho C, Coulouris G, Avagyan V, Ma N, Papadopoulos J, Bealer K, et al. BLAST+: architecture and applications. BMC bioinformatics. 2009;10(1):421.

59. Waterhouse AM, Procter JB, Martin DM, Clamp M, Barton GJ. Jalview Version 2—a multiple sequence alignment editor and analysis workbench. Bioinformatics. 2009;25(9):1189–91.

60. Crooks GE, Hon G, Chandonia J-M, Brenner SE. WebLogo: A Sequence Logo Generator. Genome Research. 2004;14(6):1188–90. doi: 10.1101/gr.849004. PubMed PMID: PMC419797.

61. Davis JJ, Olsen GJ. Modal codon usage: assessing the typical codon usage of a genome. Molecular biology and evolution. 2009;27(4):800–10.

62. Davis JJ, Olsen GJ. Characterizing the native codon usages of a genome: an axis projection approach. Molecular biology and evolution. 2010;28(1):211–21.

63. Ranieri ML, Shi C, Switt AIM, Den Bakker HC, Wiedmann M. Comparison of typing methods with a new procedure based on sequence characterization for *Salmonella* serovar prediction. Journal of clinical microbiology. 2013;51(6):1786–97.

64. Zhu L, Olsen RJ, Nasser W, Beres SB, Vuopio J, Kristinsson KG, et al. A molecular trigger for intercontinental epidemics of group A *Streptococcus*. The Journal of clinical investigation. 2015;125(9):3545–59.

65. Nasser W, Beres SB, Olsen RJ, Dean MA, Rice KA, Long SW, et al. Evolutionary pathway to increased virulence and epidemic group A *Streptococcus* disease derived from 3,615 genome sequences. Proceedings of the National Academy of Sciences. 2014;111(17):E1768–E76.

66. Yoshida H, Kojima T, Yamagishi J-i, Nakamura S. Quinolone-resistant mutations of the gyrA gene of *Escherichia coli*. Molecular and General Genetics MGG. 1988;211(1):1–7.

67. Yoshida H, Bogaki M, Nakamura M, Nakamura S. Quinolone resistance-determining region in the DNA gyrase gyrA gene of *Escherichia coli*. Antimicrobial agents and chemotherapy. 1990;34(6):1271–2.

68. Tyson GH, Zhao S, Li C, Ayers S, Sabo JL, Lam C, et al. Establishing genotypic cutoff values to measure antimicrobial resistance in *Salmonella*. Antimicrobial agents and chemotherapy. 2017;61(3):e02140–16.

69. Kohanski MA, Dwyer DJ, Hayete B, Lawrence CA, Collins JJ. A common mechanism of cellular death induced by bactericidal antibiotics. Cell. 2007;130(5):797–810.

70. Foti JJ, Devadoss B, Winkler JA, Collins JJ, Walker GC. Oxidation of the guanine nucleotide pool underlies cell death by bactericidal antibiotics. Science. 2012;336(6079):315–9.

